# Understanding The Role of Heparinoids on the SARS-CoV-2 Spike Protein through Molecular Dynamics Simulations

**DOI:** 10.1101/2022.07.05.498807

**Authors:** Ludovico Pipitò, Christopher A. Reynolds, Giuseppe Deganutti

## Abstract

The pandemic caused by severe acute respiratory syndrome coronavirus 2 (SARS-CoV-2) continues to pose a threat, with an estimated number of deaths exceeding 5 million. SARS-CoV-2 entry into the cell is mediated by its transmembrane spike glycoprotein (S protein), and the angiotensin-converting enzyme 2 (ACE2) receptor on the human cell surface. The extracellular heparan sulphate (EcHS) enhances the S protein binding through a mechanism that is still unknown. Surprisingly, low molecular weight heparin (LMWH) and HS in the disaccharide form (dHS) hinder the S protein binding to ACE2, despite the similarity with EcHS. We investigated the molecular mechanism behind this inhibition through molecular dynamics (MD) simulations to understand the interaction pattern of the heparinoids with S protein and ACE2 receptor.

## Introduction

The severe acute respiratory syndrome coronavirus 2 (SARS-CoV-2) spike protein (S protein) has a strong affinity for the human angiotensin-converting enzyme 2 (ACE2) receptor, a type 1 transmembrane protein responsible for the extracellular conversion of the angiotensin hormone into angiotensin II (1). The S protein (**Figure 1A**) is a highly glycosylated, conserved trimeric structure amongst the *coronaviridae* family. Each S protein monomer can be divided into two main domains, S1 and S2 (2). While S1 recognizes ACE2, S2 perforates the membrane of the host cell to transfer the genetic material into the cytoplasm. The receptor-binding domain (RBD) of S1 (residues R319-F541) is responsible for the molecular recognition of ACE2 (**Figure 1B**) and for triggering conformational changes that initiate the infectious mechanism.

**Figure 1.**
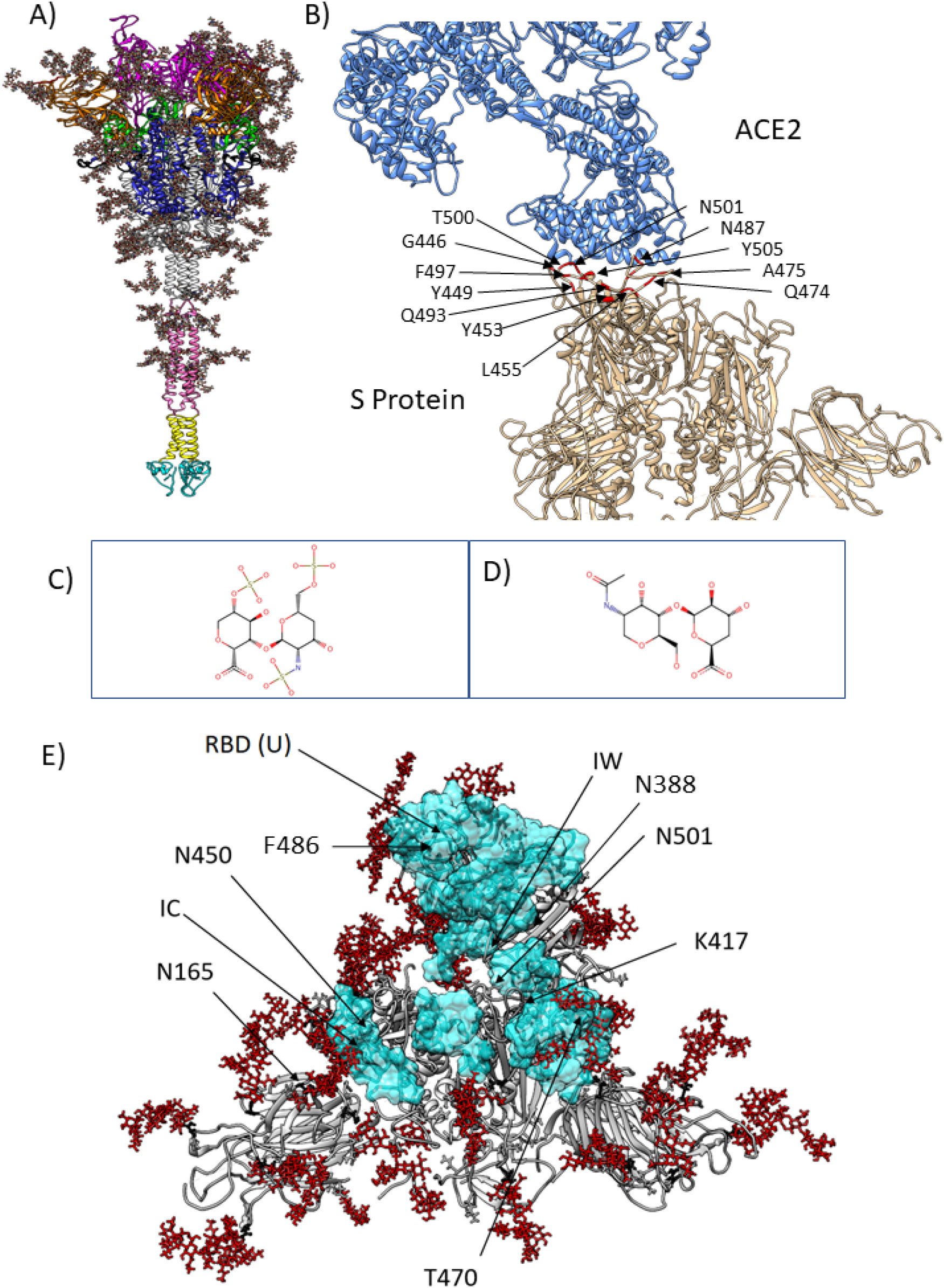
The S protein with its domains and docking results. **A)** The fully glycosylated S protein with each domain coloured as follows: signal peptide (SP) (red), N-terminal domain (NTD) (orange), receptor binding domain (RBD) (magenta), C-terminal domain (CTD) (green), S1/S2 cleavage site (Black), fusion peptide (FP) (Blue), heptad repeat 1 (HR1) (light grey), heptad repeat 2 (HR2) (pink), transmembrane domain (TM) (yellow), and cytosolic domain (CD) (cyan). Glycans are depicted in stick. (3). **B)** S1 (grey) - ACE2 (blue) complex; RBD “up” residues (red) close to ACE2 are responsible for the molecular recognition. **C)** Heparin most representative disaccharide unit (dHP), consisting of alpha-L-iduronate 2-O-sulfate and glucosamine 2,6-disulfate. **D)** The dHS model used for simulations (glucuronic acid and N-acetylglucosamine). **E)** Top view of molecular docking results. dHS and dHP (cyan surface) were predicted to bind a large area of the S protein (grey ribbon) corresponding to the internal wall (IW) of RBD “up” from K417 to N501 and the internal channel (IC) formed between residues N450 and T470 between NTD and the RDB “down”. HS is encompassed between the glycan residues (red stick representation) on N165 and the RBD down region between N450 and N501 and between the gaps between RBD and NTD.

Several S protein structures in the metastable pre-fusion state (**Figure 1 A**) have been solved through cryo-electron microscopy (cryo-EM) (4)(5), allowing the distinction of two different RBD configurations, named “up” and “down” (5), the former responsible for binding to ACE2. A set of 13 RBD residues (T402, R439, Y436, N440, Y455, N473, Y475, F486, G488, Y491, Q493, Q498, N501) is involved in interactions with ACE2 (6) and constitutes the receptor binding motif (RBM) on the apical portion of RBD (**Figure 1B**) (7).

Glycosaminoglycans (GAGs), such as extracellular heparin (EcHP) and heparan sulphate (EcHS), play a crucial role in regulating the immune response by regulating cell adhesion, tuning cytokine, and chemokine function, and mediating inflammatory reactions (8,9), through HS-binding motifs (HSBM). EcHP, also produced by basophils and mast cells (10), is constituted by highly sulphated repeating units of 1-4 pyranosyl uronic acid and 2-amino-2-deoxy glucopyranose (**Figure 1C**) (glucosamine) and it is known for its major role as an anticoagulant when formulated in low molecular weight (LMWH). EcHP is characterized by a wide structural heterogeneity and high negative charge due to the presence of numerous sulphate groups per disaccharide units of O-sulphonated D-glucosamine and O-sulphonated hexuronic acid (**Figure 1D**) (11). The high level of sulphonation makes the EcHP formal charge higher than EcHS (9) and is responsible for LMWH pharmacodynamics properties on the coagulation cascade by binding to the antithrombin (12). EcHS is diffusely expressed on the cellular surface in the form of high molecular weight heparan sulphate proteoglycans (13). EcHS has various roles as immune regulator (8), anticoagulant (14), and reportedly assists microbial and viral infection acting as molecular adhesion receptors in human immunodeficiency virus (HIV). The role of EcHS in assisting viral infections (15) is carried out both by promoting virus-host cell adhesion and by acting as a “glue” between viral proteins and host cell receptors.

EcHP has been proposed to play a role in promoting SARS-Cov-2 infection (16,17), probably by inducing conformational changes upon binding (18) through interactions on the S1/S2 cleavage site (19). The intermediary role of EcHS between the furin cleavage site and GAG was also suggested by Schuurs and co-workers (20), who proposed the involvement of GAGs in favouring the membrane fusion mechanism. Clausen and collaborators (21) proposed EcHP to allosterically facilitating interactions between S protein and ACE2 and provided a preliminary model of the RBD regions possibly implied in the recognition of both EcHS and EcHP. Other studies suggested the intriguing hypothesis that EcHP and low molecular weight heparins (LMWH) may function as antagonists (22,23) of ACE2 binding by competing for the EcHS binding site on S protein (24).

We investigated the possible binding differences between heparinoids (13) on both S protein and ACE2 receptor by using a mixed molecular dynamics (MixMD) approach (25), to expand the model of binding to S protein and clarify whether the sulphated groups, abundant on EcHP, could play a role in the SARS-CoV-2 binding to ACE2.

## Methods

### Structure Preparation and Force Field Settings

All systems were prepared using the CHARMM36 (26,27)/CGenFF 3.0.1 (28,29) force field combination. To speed up the simulations, only the apical portion of the fully glycosylated S protein was kept from the spike protein model 6VSB_1_2_1 available at (https://charmm-gui.org/?doc=archive&lib=covid19) (30). Such regions (residues 1-1003) included the S1 and S2 domains SP (1 - 13), NTD (14-305), RBD (306-541), CTD (542 - 652), S1 / S2 (653 - 686), and FP (687-911). The dHP and dHS (**Figure 1C, 1D**) were designed with the VMD Molefacture plugin (https://www.ks.uiuc.edu/Research/vmd/plugins/molefacture/).

ACE2 peptidase domain dimer (ACE2-PD) was taken from the PDB entry 6M17 (residues I21-L731) (31). Hydrogen atoms were added to the S protein and ACE2 by propka (32) at a simulated pH of 7.0, while disulphide bonds were identified by HTMD (33), visually inspected, and patched manually through VMD (34) as previously reported (35). The resulting structure were energy minimized using the conjugate gradient algorithm by ACEMD (33).

### Molecular Docking

Multiple docking experiments were performed using Autodock Vina (36) on the apical portion of the fully glycosylated S protein S1 domain. As a reference for docking, the centre of the grid box was located on residue C432 of each monomer in the “up” and “down” state **(Figure 1E)**. The docking was repeated six times by moving the box +5 Å on the z-axis on the RBD “up” to include residue Q493-Y505. Finally, a blind docking was performed on the overall S protein (**Table S1**).

### MixMD system preparation

To map possible S protein and ACE2 sites interacting with HS and HP, each probe was simulated individually using a MixMD approach (37). In total, four systems were prepared using dimeric HP or dimeric HS through Packmol (38) (**Table S2**). A minimum distance of 4 Å was set to avoid clashes and secure an ordered placement of the molecules within a 150 Å x150 Å x150 Å box (39). TIP3P water molecules (40) were added to the simulation box considering a 15 Å water padding by Solvate plugin 1.5 (http://www.ks.uiuc.edu/Research/vmd/plugins/solvate/). The charge neutrality of the system was achieved by adding Na^+^ /Cl^−^ to the concentration of 0.150 M using Autoionize plugin 1.3 (http://www.ks.uiuc.edu/Research/vmd/plugins/autoionize/). ACEMD was used for both equilibration and MD production. The energy of the systems was reduced through 2000 conjugate-gradient minimization (CG) steps to eliminate possible clashes and optimize atomic distances. Equilibration was reached in isothermal-isobaric conditions (NPT) using the Berendsen barostat (41) (target pressure 1 atm) and the Langevin thermostat (42) (target temperature 310 K) during a 4 ns simulation (integration time step 2 fs). During the equilibration, a positional restraint of 1 kcal/ mol Å^2^ was applied only on the alpha carbons at the base of the S protein, where the structure was cut from the rest of the protein, leaving the glycans and the three RBDs free to move. Two 500 ns productive trajectories were produced with an integration time step of 4 fs, using the hydrogen mass repartition (HMR) (43), in the canonical ensemble (NVT), with the same positional restraints used in the equilibration. The cut-off distance for electrostatic interactions was set at 9 Å, with a switching function applied beyond 7.5 Å. Long-range Coulomb interactions were handled using the particle mesh Ewald summation method (PME) (44) with default ACEMD settings. In summary, two trajectories were produced for each ACE2-HP, two for ACE2-HS, two S protein-HP, and two S protein-HS, reassigning the atomic velocities on each replica, for a total of 1000 ns for each system.

### MD trajectories analysis

MD Trajectories were merged using MDtraj (45). The root-mean-square deviation (RMSD) and root mean square fluctuation (RMSF) analysis were computed using VMD (34). Ligand-protein contacts, including hydrogen bonds, were detected using GetContacts (https://getcontacts.github.io), with a threshold distance and angle of 3.5 Å and 120°, respectively. Contacts and hydrogen bonds were expressed as occupancy (% of total MD frames) over the two merged replicas for each system. Molecular graphics images were produced using UCSF Chimera (46). Volumetric maps were computed using the VMD VolMap plugin with a space grid of 0.25 A, averaging all frames (https://www.ks.uiuc.edu/Research/vmd/vmd-1.9.1/ug/node153.html).

## Results

Low molecular weight heparan sulphates (LMWHS) have been proposed as possible antagonists of the ACE2 binding to the S protein (22) thanks to the high degree of sulfonation (17). We tried to clarify the potential role of sulphate groups on S protein binding to better understand this mechanism, focusing on crucial S protein regions involved in ACE2 molecular recognition (7).

### Molecular Docking Suggests Broad Distribution of HP and HS Interaction Sites

Molecular docking of disaccharide HP (dHP) and HS (dHS) was performed to replicate the data obtained by Clausen *et al*. (21). In Clausen’s study, tetra-saccharide heparin (dp4) units were docked against RBD in both “up” and “down” states, suggesting interactions with residues R346, R355, K444, and R466, while F347, S349, N354, G447, Y449, and Y451 were suggested favouring the S protein - ACE2 alignment. Our docking agreed with the previous results, showing a homogeneous distribution of poses on S protein. A broader distribution was predicted on the “up” conformation, in correspondence of residues N388 - F486 (**Figure 1E**). Docking on RBDs “down” indicated a lesser engagement, with HS and HP molecules predicted on a smaller surface, close to residues K150 and T470 into the cleft between NTD and RBD (**Figure 1E**). Additionally, HS was also suggested to bind the RBD, responsible for the molecular recognition with ACE2, at the level of residues K417-T500 but with the lowest docking score among all poses. These initial results suggests that the disaccharides are likely to lodge into niches at the base of RBD, rather than engaging with the ACE2 binding site directly

### MD simulation elucidates the flexibility of the S protein and its glycoshield

Although molecular docking predicted a large area of the S protein potentially involved in interactions with dHS and dHP, it did not consider the protein flexibility and explicit solvent. To overcome these intrinsic limits, we performed MixMD simulations to sample S protein’s sites potentially interacting with HS and HP, in a flexible and fully hydrated environment (**Video S1, Video S2**). During the MixMD replicas, the RBDs “down” remained stable (**Figure S1, Figure S2**) due to the presence of salt bridges between K378-E988 and K386-D985 in the S2 domain, and E516-K202 in the N-terminal domain (NTD), in accordance with Gur *et al* (47). The RBD “up” showed overall higher mobility (**Figure S1, Figure S2**) but moderate flexibility in the loop between F485 and Y505 and high flexibility in the I468-Q493 region. The glycans bound to N165 intercalated between N427 and E465 at the base of the RBD, preventing it to fold back to the “down” conformation (48). Overall, glycans displayed the highest mobility amongst all the residue in the system (**Figure S3**). Sugars bound to N165, N234, and N343 restricted the access to the apical portion of S1, while the glycans linked to N165 locked RBD in the “up” state, in agreement with Casalino and co-workers (49). These results indicate that the S protein dynamics, in conjunction with a sweeping motion of the glycans, might restrict the access to the lateral wall of RBD (residues R346, N354-R357) and to the niche between RBDs “up” and the “down”, as well as on residues R403-R408 at the centre of the S1 domain. Areas across NTD and close to RBD “up” offered better chances to engage dHS and dHP in productive contacts due to the lower shielding of glycans.

### Simulations identified similarities in HP and HS interactions with S protein

MixMD simulations of dHS and dHP against the glycosylated S protein indicated similar distributed areas involved, including NTD and both RBDs “up” and “down”. Both dHP and dHS engaged the S protein in persistent contacts with areas not covered by the glycans’ sweeping motion: **i)** RBD residues R346, N354-K356, R408, K458, S477, and to a lesser extent to N501-G504, on the RBM; **ii)** residues R158, N164, I231, R237, H245-R246 on the NTD, and **iii)** residues F4, Q14, N17, S98, K147, R158, K182-G183, H245, and G252 on the external projections of NTD (**Figure 2, Figure S4**) indicating competition between the two ligands for multiple sites.

**Figure 2.**
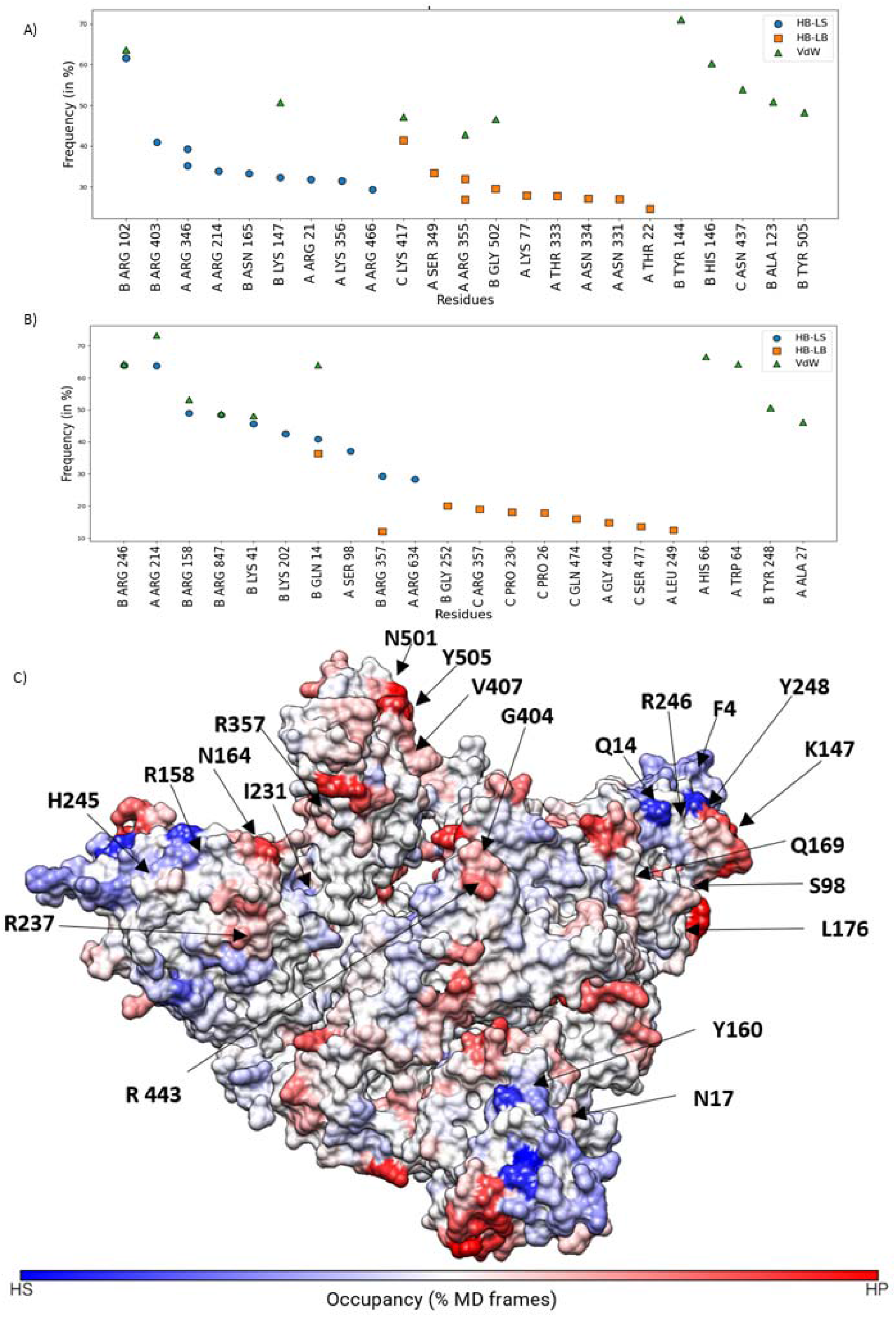
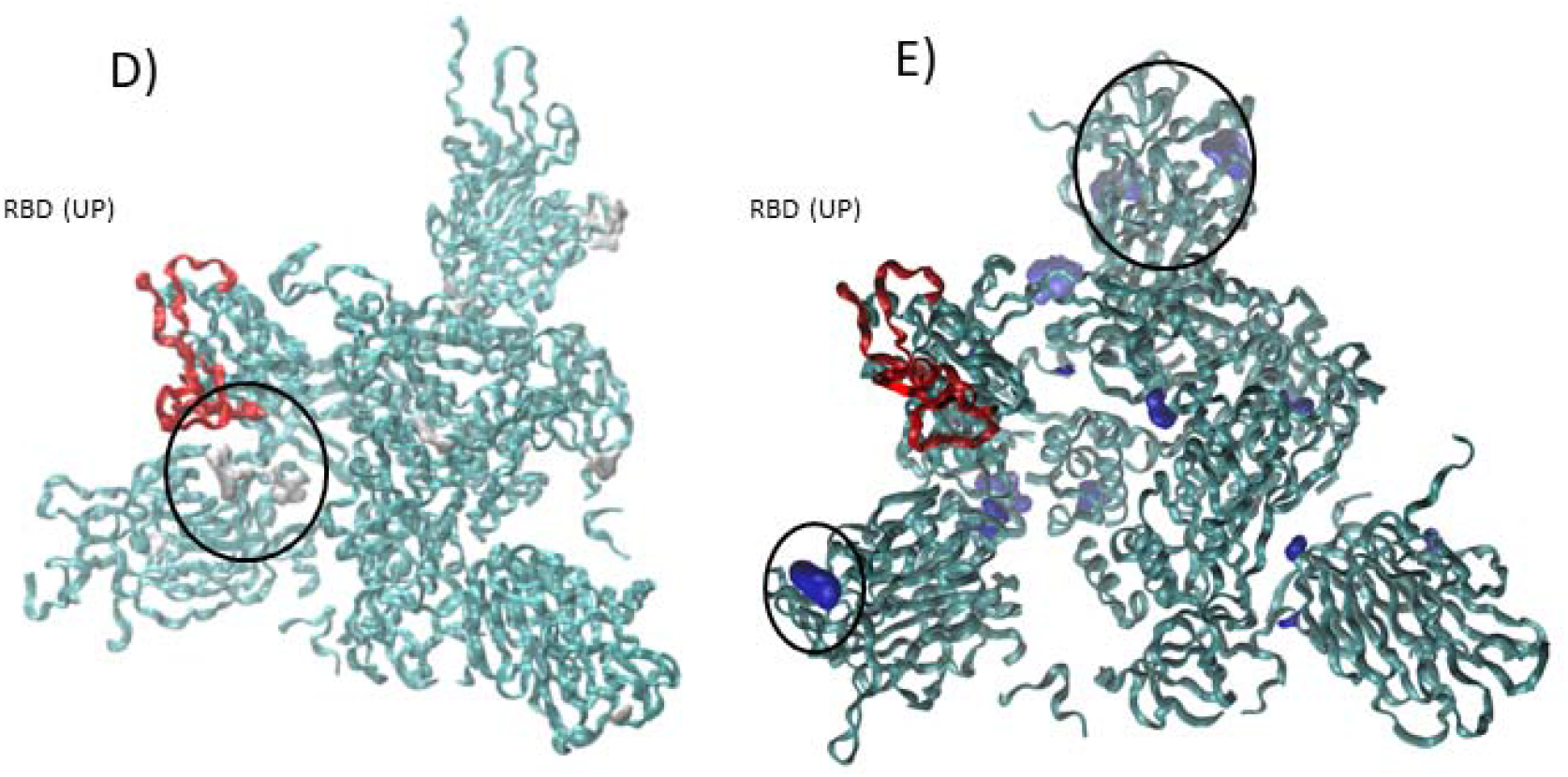
Interactions between disaccharide HS or HP and the S1 domain. **A)** Residues most interacting with dHP, ranked according to the frequency of interactions during the MixMD simulations. G502 and G504, which are part of the RBM, showed multiple contacts with dHP; **B)** residues most interacting with dHS, ranked according to the frequency of interactions during the MixMD simulations. “A” indicates the RBD “up”, while “B” and “C” indicate the RBD down; HB-LS: hydrogen bonds involving S protein sidechains; HB-LB: hydrogen bonds involving the S protein backbone; VdW: van der Waals interactions. **C)** Differences in S protein contacts with dHS or dHP, plotted on the S protein (ribbon representation) and coloured according to the occupancy prevalence (% MD frames). The polar residues R158, N164, R357, and V407 made more contacts with dHP, while F4, Q14, N17, Q169, Y248 engaged more dHS, at the “corners” of S protein, in agreement with Clausen’s results. **D)** Volumetric density map of dHP (grey, isovalue 20%) showing the averaged atomic positions for the ligand across the simulation box. **E)** Volumetric density map of dHS (blue, isovalue 20%); The circles in both images indicated the spots where the ligand persistence remained constant regardless of the set isovalue around NTD, SP domains, located at the base of S1. The red ribbon highlights the RBD in the up state for on chain A of the trimer.

Many persistent contacts occurred between dHP and most of the S protein surface, specifically on residues R355-R357, C166, R466, and R346, and the inner portion of the two RBD “down” on residues N501-Y505, R408 (**Figure 2C**). Residues R355-R357, R408, K529, and G502-G504 were engaged by dHP in both RBD “up” or “down” conformations. Notably, none of these residues is part of the binding site for ACE2, except for N501-Y505. dHP engaged the R403-R408 region in both two “down” RBDs as well in the centre of the trimer, involving G502 and G504 on every chain. dHP bound more prevalent on residues N164, I231, G404, R408, G502-G504, in both RBD “up” and “down”, and R355-R357, V407, and R408 of the RBD “up” state. dHP formed more persistent interactions than dHS with R354-K356, probably due to negative charge of its sulphate groups.

Overall, dHP and dHS contacts (**Figure 2C**) substantially overlapped, suggesting no preference of the S protein against any heparinoid. However, the volumetric density maps highlighted some differences between HP and HS (**Figure 2D, 2E**). The latter, indeed, stationed by the NTD apical region close to the S protein base (residues S98-K147, R237, R246-Y248, R357) and by the RBD “up” residues R355-R357, K378, G404. Residues V407 S477 and Q474, and the surrounding areas at the centre of the trimer (**Figure 2C**) were instead equally engaged by HP and HS. The volumetric density maps indicate higher occupancy of the dHS on the corners of NTD and in the proximity of the signal peptide (SP.) (**Figure 2D, 2E**).

In summary, both dHS and dHP mapped similar hotspots on the S protein, highlighting three main sites of interaction: regions delimited by the R346-R357 facing the adjacent NTD residues N165-S172, residues N405, R408, G502-G504 on RBD “up”, and residues P26, K97, S98, K182, and P251 at the corner of each NTD, suggesting no marked preference of the S protein for any of the two heparinoids. Altogether, our simulations indicated very low occupancies near the ACE2 binding site, suggesting low affinity for dHS or dHP. The protruding portion of the NTD may represent a possible engagement point for EcHS, providing support in the alignment of the S protein and ACE2 (**Figure 3F**), while EcHP could support the RBD “up” conformation without interfering with the ACE2 interaction. The contacts between G502 and Y505 would provide interactions sites for EcHP to support the opening of RBD in the “up” conformation, in agreement with Clausen’s results.

**Figure 3.**
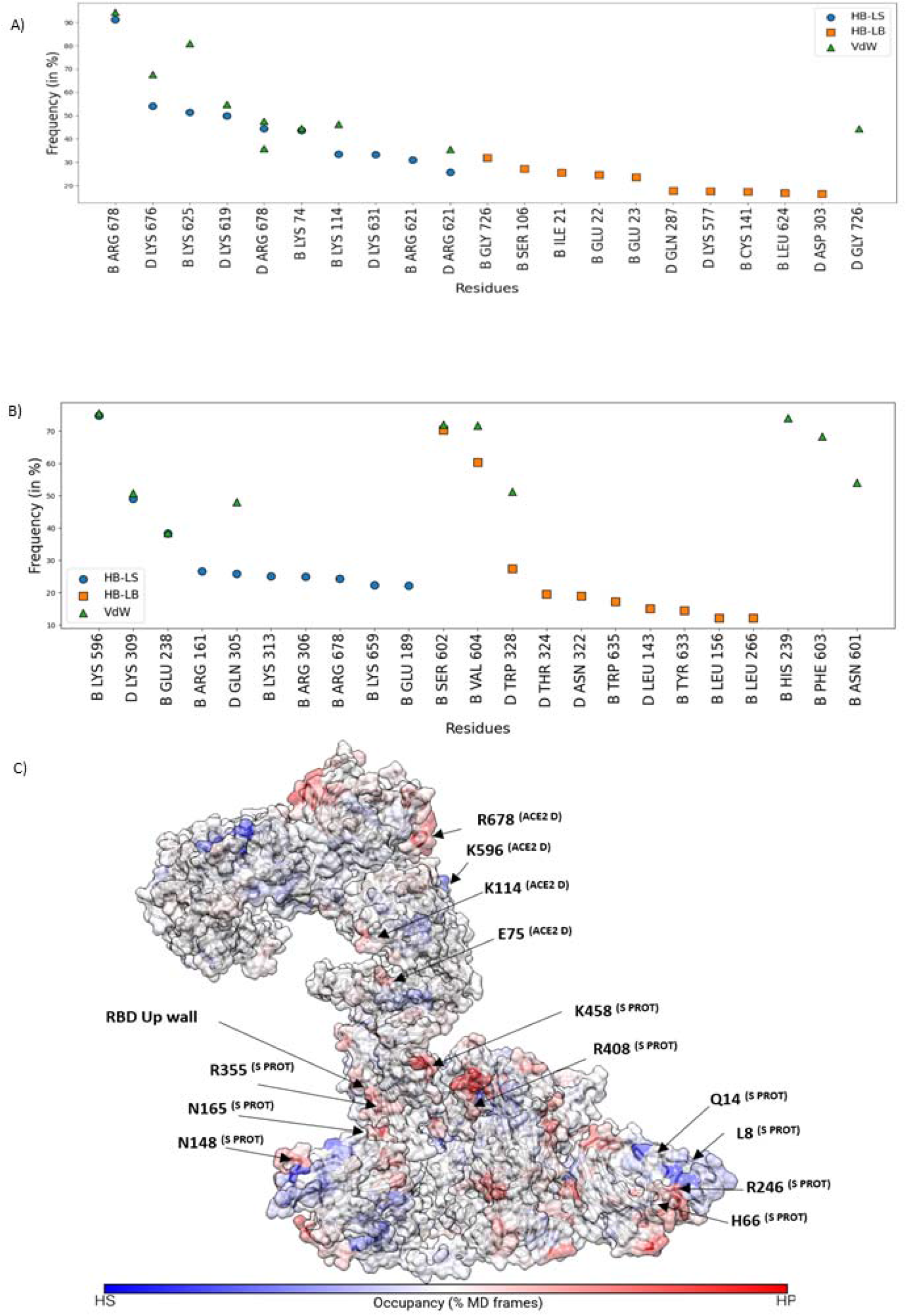
Interactions between HS or HS and ACE2. **A)** Contact frequency between ACE2 residues with dHP, ranked according to the frequency of interactions during the MixMD simulations. Chain B and D are respectively the two extracellular PD domains with chain D being the one depicted on the right. Persistent interactions involved predominantly residues located on the extracellular domain of ACE2 in proximity with the cellular membrane, especially around R678, K676, K625. The letters on the table indicate the chain where the residue is located. HB-LS: hydrogen bonds involving S protein sidechains; HB-LB: hydrogen bonds involving the S protein backbone; vdW: van der Waals interactions. **B)** Table representation of most interacting ACE2 residues with dHS, ranked according to the frequency of interactions during the MixMD simulations. Persistent interactions were produced on the PD domain on residue K 596 and the surrounding area, K309 on the front helix, and K114 located inside the PD. **C)** dHP and dHS atomic contacts plotted on S protein-ACE2 complex and coloured according to the occupancy differences.

### Disaccharide HP and HS interaction sites on ACE2

We focused on ACE2 to pinpoint possible interaction sites for EcHP and EcHS, involved in the enhanced binding between ACE2 and S protein(50,51). During MixMD simulations, ACE2 pincer-shaped domains underwent an outward opening movement (**Video S3, Video S4**) and exhibited flexibility at the PD domain level on chain D (**Figure S5, Figure S6**). dHP and dHS interacted on three main sites of the PD **Figure S7**).: **i)** residues I21, Q22, K74, K114, and R306-K313 on the external portion of the pincer formed by the two units, **ii)** the junction area at residue Q140, G287, D303, K577, and **iii)** residues K619, R621, K625, K631, K676, R678, K702, G726, which delimit a ring around the PD base (**Figure 3A**).

Overall, the two disaccharides tent to engage similar residues around the extracellular domain of ACE2, on the sides of PD, and across the helical bundle, although the higher sulfonation of dHP increased electrostatic interactions with lysine and arginine residues. From a mechanistic perspective, interactions are predominant at the base of ACE2, around residues K625, K676-R678, and G726. EcHP could bind in this region without blocking ACE2 access and at the same time provide the S protein with a closer contact to the host’s membrane. dHS formed more persistent interactions than dHP with the helix bundle identified by residues K309-W328, suggesting low competition between the two ligands in that region. dHS engaged ACE2 in three distinct zones: **i)** on residues Q305-R306, K309, K313, T324, W328 in the helical bundle, **ii)** on residues E238, K596, S602 in the outer wall of the monomers, **iii)** at the base ACE2 extracellular domain, on residues W635-N636, K659, and R678 (**Figure 3C, Figure S7, Figure S8**). Like dHP, regions at the base and outside the PD formed more stable contacts throughout the simulations. dHS interacted with residues N148, N165, R355, R408, and K458, and on the internal walls delimited by RBD “down” residues R408, K444, and G502. The most persistent contacts between dHP and ACE2 occurred at the base of the receptor. dHS probes identified specific sites on the NTD around residues F4, L8, Q14, H66, S98, K147.

In summary, simulations indicated that EcHP and EcHS would not interact with the S protein binding site for ACE2 and suggested that long chained heparinoids bound to the S protein can extend toward the base of the ACE2, providing an additional anchoring point to facilitate viral binding.

## Discussion

We have identified potential sites on the S protein and ACE2 receptor where EcHS and EcHP could bind and compete. Our results indicates that both heparinoids can linger between the RBD external wall, around R355, and the adjacent NTD close to N165. dHS formed more contacts at the corners of each S protein monomer NTD domain, including a portion of the SP. Strong similarities between the interactions heatmaps of dHS and dHP indicate that SARS-CoV-2 could exploit both EcHP and EcHS, regardless of their sulfonation state, to anchor itself in the proximity of the cellular membrane by using the proteoglycans-bound EcHS or free form EcHP with a stronger affinity for highly sulphonated disaccharides. The presence of restricted S protein areas selective for either of the two heparinoids suggests that long EcHS chain could unwind from K148, pass close to N165, and stabilise the RBD “up” in correspondence of N165 and between R355-R357 (**Figure 4**). An alternative hypothesis is that the S protein exploits both EcHP and EcHS on residues G404, R408, K444, G502-G504, to favour the initial RBD transition to the “up” state with EcHP supporting the transition between the states in a more efficient manner. Hence, the role of heparinoids would be to favour, together with the glycan on N165, the opening and stabilization of RBD towards the “up” state. S protein would be evolutionarily designed to exploit the molecular similarities between EcHP and EcHS to approach the membrane through the residues R346, R355-R377, K444 for activation and alignment. This hypothesis sees EcHS as a landing “hook” for the S protein NTD, favouring the correct orientation of RBD toward ACE2, as indicated by the numerous contacts on multiple residues on NTD. In this scenario, unbound EcHP could additionally stabilize and open the RBD while anchoring S protein to ACE2, as indicated by the contacts on both the S protein and the extracellular portion of ACE2, suggesting a synergic effect of EcHS and EcHP, in agreement with Clausen’s results. Long EcHS chains projected towards the extracellular lumen could intercept S protein through electrostatic interaction between sulphated groups and positively charged amino acids exposed on S1 and NTD, facilitating the ACE2 binding in agreement with Tandon’s group experiments.

**Figure 4.**
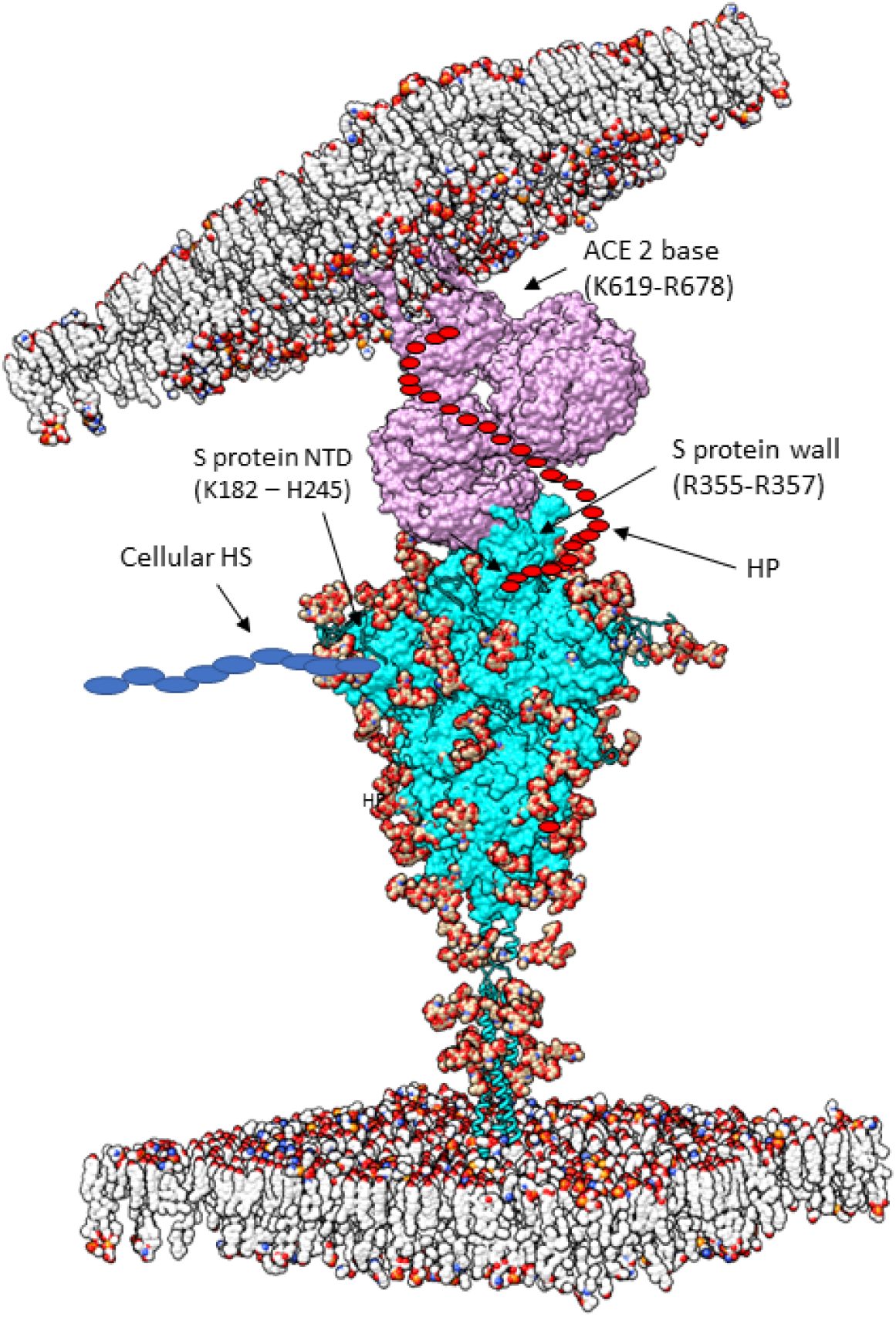
Proposed interaction model for EcHS and EcHP on the S protein-ACE2 complex. The presence of EcHS in the glycocalyx on the host’s extracellular matrix could be exploited by SARS-2 to better anchor and align the virion to the host cell, without interfering with the RBM. Long EcHS chains could also act as three-point adhesives, enhancing the SARS-2 infective mechanism. HP could lock RBD in the “up” state without affecting the molecular recognition with ACE2.

It is reasonable to assume that the inhibition experimentally displayed by dHP and dHS is due to the ability of short chains to saturate all the binding sites for EcHS and EcHP onto S protein and ACE2 surface, preventing longer EcHP chains to bind and facilitate the conformational shift of RBD from “down” to “up”, or hindering the alignment with ACE2.

## Conclusion

We performed MixMD simulations of dHP and dHS against the glycosylated S protein and the ACE2 receptor, identifying common zones and specific sites where each ligand predominantly engages. Our results support the scenario in which shorter heparinoid could saturate S protein binding spots on the NTD and RBD and interfere with the S protein alignment and stabilization exerted by the extracellular matrix in proximity of the membrane. The abundance of S protein sites affine to both dHS and dHP also suggest that the saturation of S protein sites by short chain HS and HP could prevent EcHS-dependant viral binding and the correct RDB opening.

