## Supplementary material for "Understanding The Role of Heparinoids on the SARS-CoV-2 Spike Protein through Molecular Dynamics Simulations": Table S1

### Table of content

**Video files**

**Table S1 Initial Molecular Docking input**

**Table S2 MD Simulation Summary**

**Figure S1 Spike protein RMSF and RMSD during MixMD simulations in the presence of HP.**

**Figure S2 Spike protein RMSF and RMSD during MixMD simulations in the presence of HS.**

**Figure S3. RMSF and RMSD of glycan residues on the S protein.**

**Figure S4. S Contacts between S protein and HP and HS.**

**Figure S5. ACE2 RMSF and RMSD during MixMD simulations in the presence of HP.**

**Figure S6. ACE2 RMSF and RMSD during MixMD simulations in the presence of HS.**

**Figure S7. S Contacts between ACE2 and HP (red) and HS (blue).**

**Figure S8. Volumetric maps (ISOVALUE = 20%) of HP (top) and HS (bottom)**

**Video S1.**

**Left hand side.** S1 domain with RBD up colored in red in presence of 100 HP units. dHP units make frequent contacts at the base of RBD “up”, specifically between R355-R357 and in proximity of G502-G504 on the RBD “down”. Furthermore, the dHP units contact NTD on H146-N148, around each corner of the structure.

**Right hand side.** S1 domain with RBD up colored in red, and density maps (20% isovalue) colored in blue. These maps identify three main areas where the density becomes relevant: at the base of RBD “up”, the center of the trimer, between the two adjacent RBD “down” and between RBD “down” and NTD. The niche between RBD “up” and NTD suggests that long EcHP chains could lodge in between to support the stabilization of the “up” conformation of RBD, by preventing the downward folding motion, in conjunction with glycans sweeping motion (not shown in the video for clarity).

**Video S2.**

**Left hand side.** S1 domain with RBD up colored in red in presence of 100 HS units. dHP units make frequent contacts at the base of RBD “up”, specifically between R355-R357 and in

proximity of G502-G504 on the RBD “down”. Furthermore, the dHP units contact NTD on H146-N148, around each corner of the structure.
**Right hand side.** S1 domain with RBD up colored in red, and density maps (20% isovalue) colored in blue. Different distinct areas are shown in blue, which are located on specific areas of the S1 domain, especially around the NTDs, suggesting a specific preference from dHS to the peripheral spots across the S1 domain. This would suggest that the spike protein could specifically exploit the extracellular heparan sulphate, bound to proteoglycans on the membrane protein to latch on it and facilitate the proper RBD – ACE2 alignment. It is important to notice how RBD has no dHS surrounding it.

**Video S3.**
**Left hand side.** ACE2 PD domain in presence of 100 dHP units. Due to the stochastic nature of the simulation, the majority of the interactions occurred on chain B (on the right), both in the cleft on residue I21-H34, Q305-K309, W328 and at the base of the PD around residues W635-N636. The abundant dHP presence on chain B on multiple sites from the top to the bottom of the chain, in addition to the lack of a stabilizing membrane, result in a multitude of positional changes of chain B in respect to chain P. The interacting effect becomes more evident as the number of dHP units increasingly binds to chain B. The limited number of dHP units used is not able to fully cover the whole ACE2 PD, depicting the contacts only on one chain. However, due to the symmetry of the receptor, it is safe to assume a specular behavior on both chains.
**Right hand side.** ACE2 PD domain density maps (20% isovalue) colored in blue. The density map shown in blue highlight the abundance of dHP at the base of ACE2, and in few regions around the cleft. The massive presence of the dHP unit at the base allows us to bring forward the hypothesis that long EcHP chains, already engaged with the base of RBD, could potentially reach ACE2 base, possibly bridging between the two structures.

**Video S4.**
**Left hand side.** ACE2 PD domain in presence of 100 ddHS units. Less HS units do not bind on ACE2, with short-lived contacts dispersed on ACE2 PD surface. Notably the base of ACE2 around W635 represents a nice spot for dHS to temporarily make contacts. However, no dHS unit was able to make a durable contact, suggesting that EcHS is unlikely to interfere with the S protein-ACE2 interaction.
**Right hand side.** ACE2 PD domain density maps (20% isovalue) colored in blue. The density maps (iso 20%). The scarce quantity of contacts between dHS and ACE2 leaves the surface of the ACE2 PD untouched, except from an area across E238, K596, S602 on the external side of chain B. In no case, dHS poses any hindrance against RBD or the S protein, suggesting that HS role could be restricted to S protein.

**Table S1. Initial Molecular Docking input**

| HP Docking Coordinates | HS Docking Coordinates |
|---|---|
| receptor = spike_cent.pdbqt | receptor = spike_cent.pdbqt |
| ligand = hp1.pdbqt | ligand = hs1.pdbqt |
| center_x = 25 | center_x = 25 |
| center_y = 0 | center_y = 0 |
| center_z = -70 | center_z = -70 |

| | |
|---|---|
| size_x = 126 | size_x = 126 |
| size_y = 126 | size_y = 126 |
| size_z = 126 | size_z = 126 |

See attached files for individual coordinates and configurations

Autodock Vina Results:
See attached folders (spike_hp1_dockings, spike_hs1_dockings)

**Table S2. MD simulations summary.**

| **Simulation Summary** | | | | |
|---|---|---|---|---|
| **Protein PDB ID** | **Ligand (# molecules)** | **Minimization (# steps)** | **Equilibration (ts, length)** | **Simulation (ts, length)** |
| 6VSB | HS (100) | 2000 | NPT, (2fs, 10 ns) | NVT (2fs, 1000 ns) |
| 6VSB | HP (100) | 2000 | NPT, (2fs, 10 ns) | NVT (2fs, 1000 ns) |
| 6M17 | HS (100) | 2000 | NPT, (2fs, 4 ns) | NVT (4fs, 500 ns) |
| 6M17 | HP (100) | 2000 | NPT, (2fs, 4 ns) | NVT (4fs, 500 ns) |

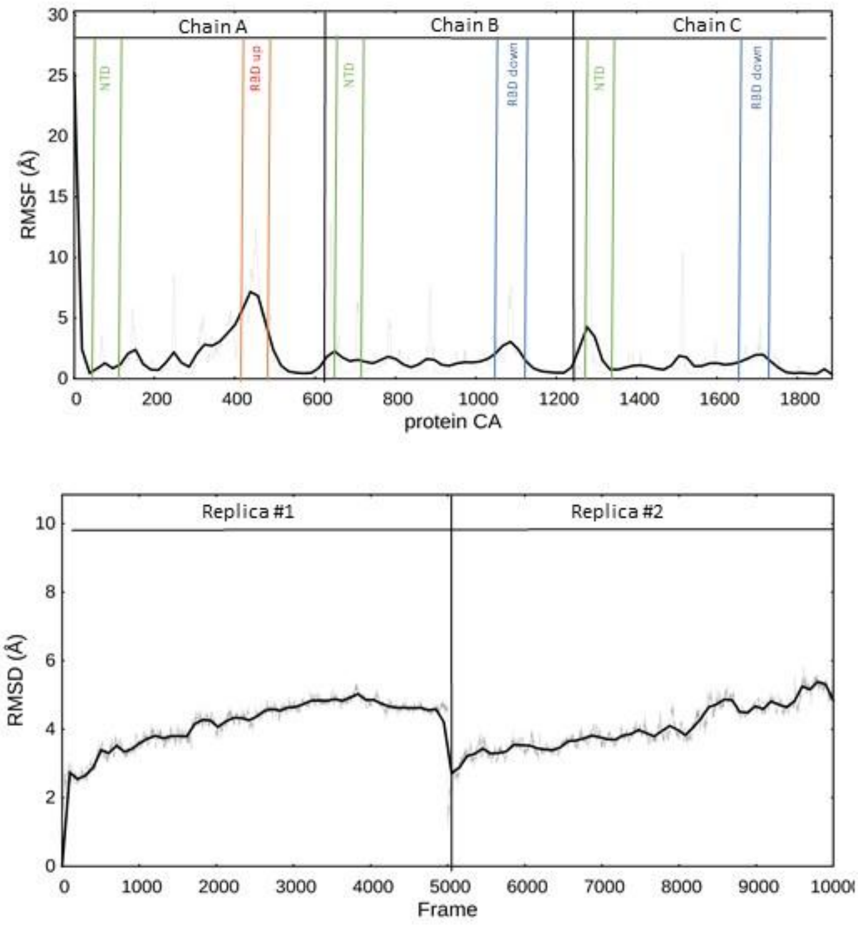


**Figure S1. Spike protein RMSF and RMSD during MixMD simulations in the presence of dHP. Results from two MixMD replicas.**

RBD displayed in the “up” conformation (chain A) displayed the highest flexibility around RBD residues R408-Y505, while the down conformations meta stable for almost the whole simulation, as indicated by the small peaks in the figure. Surprisingly, the NTD, in proximity of the SP showed a good flexibility. In our replicas, the SP adjacent to NTD on chain C showed a higher flexibility, shifting from the initial configuration and moving around its position.

The RMSD of the S1 domain gradually increased by the end of the simulation with an average RMSD value of 5 which suggests that the depth of the conformational changes that the structure can take could possibly reveal more drastic conformational changes given the removal of restraint and an increased sampling time.

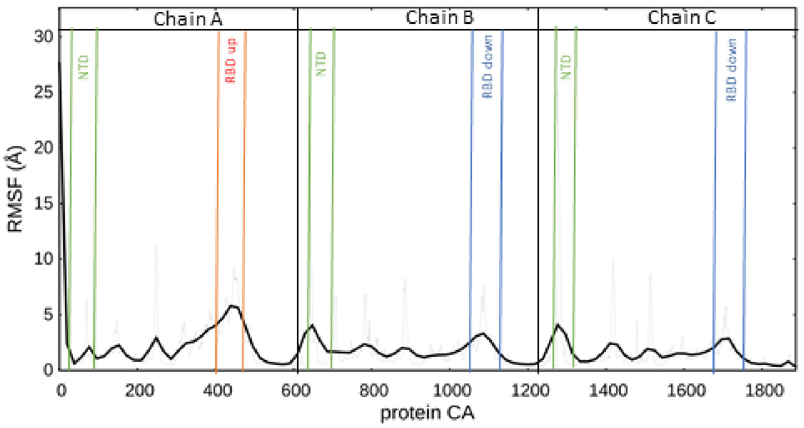


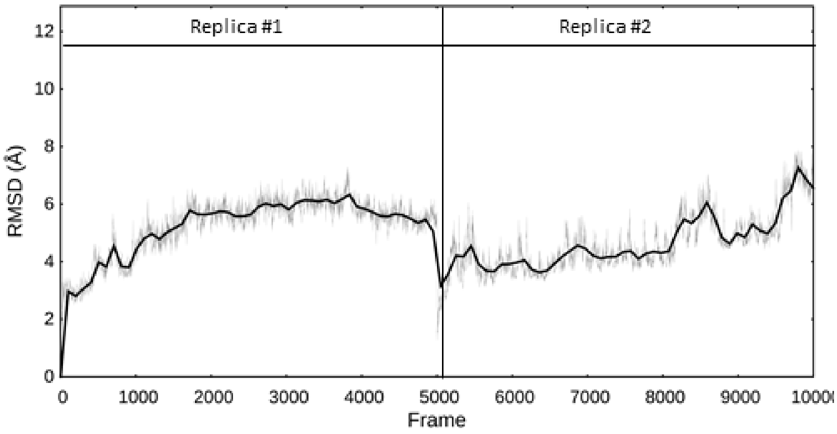


**Figure S2. Spike protein RMSF and RMSD during MixMD simulations in the presence of dHS. Results from two MixMD replicas.**

In presence of dHS, the flexibility of the whole S1 domain didn't differ much from the dHP system, except for the higher flexibility of the NTD domains. In the simulations, dHS engaged contacts with the NTD and the SP more frequently and for longer duration compared to dHP, suggesting that dHS shows a stronger occupancy for the NTD-SP corners, around residues R246-Y248, Q14, R158. However, HS was still able to interact effectively with RBD sites and across multiple areas where HP was able to interact as well, indicating an agnostic behavior of the S1 domain toward heparinoids. This hypothesis suggests an opportunistic mechanism of the whole S1 domain towards heparinoids, which could be exploited to both sustain the conformational change of RBD from "down" to "up", while using the ecHS to latch on or approach the extracellular matrix (ECM).

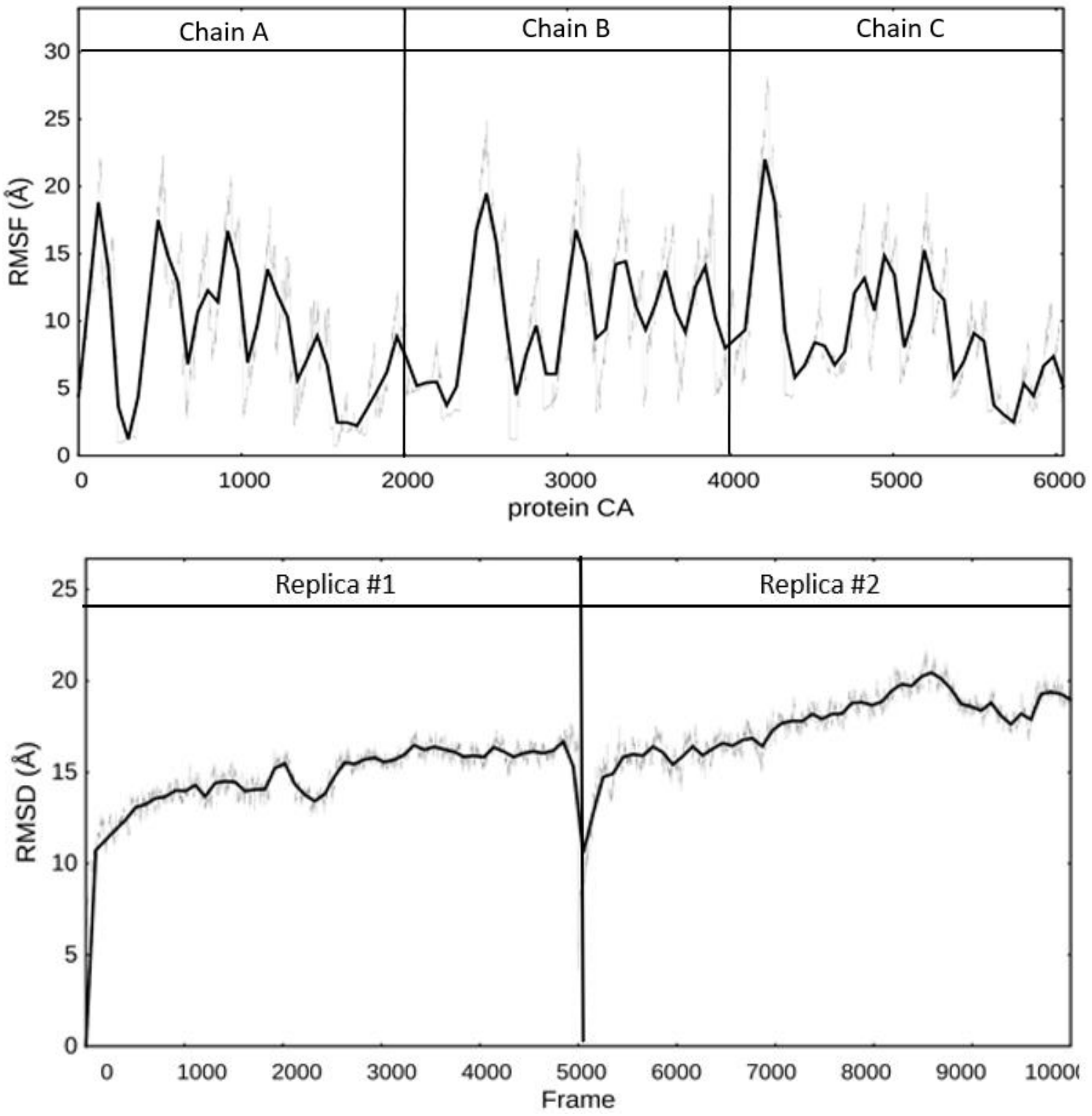


**Figure S3. RMSF and RMSD of glycan residues on the S protein. Results from two MixMD replicas.** Glycans' epitope-masking role is characterized by a protective sweeping motion which swipes the surrounding areas in a sphere of around 15 Å of radius. it follows that both the flexibility and the displacement maintain high values throughout the simulation, indicating how the flexibility of the glycans is more accentuated in the terminal portions of the glycans, indicating a "whip" behavior of the linear structure that explores the surrounding space without ever coiling on itself.

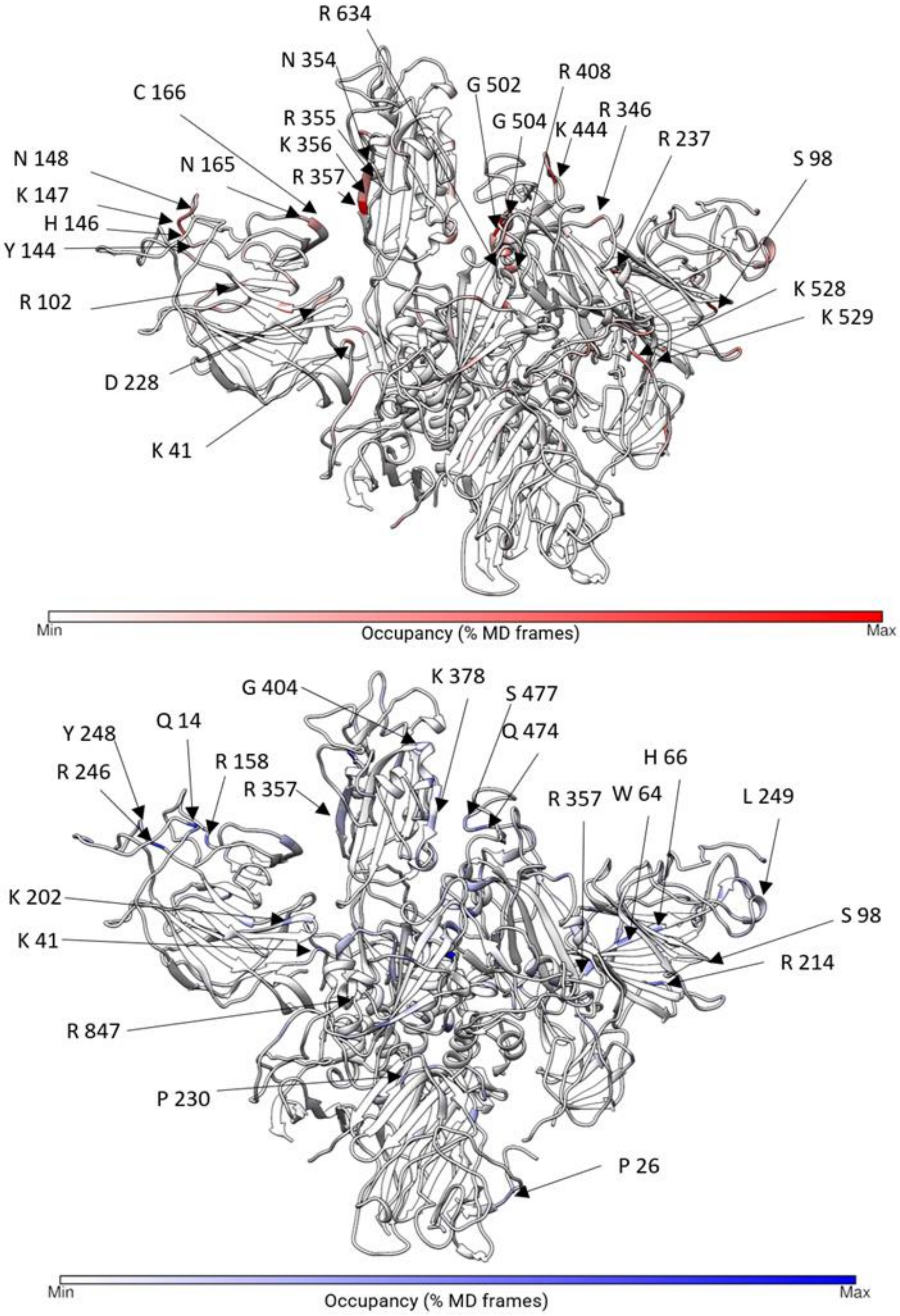


**Figure S4. S Contacts between S protein and dHP (red, top panel) and dHS (blue, bottom panel).**

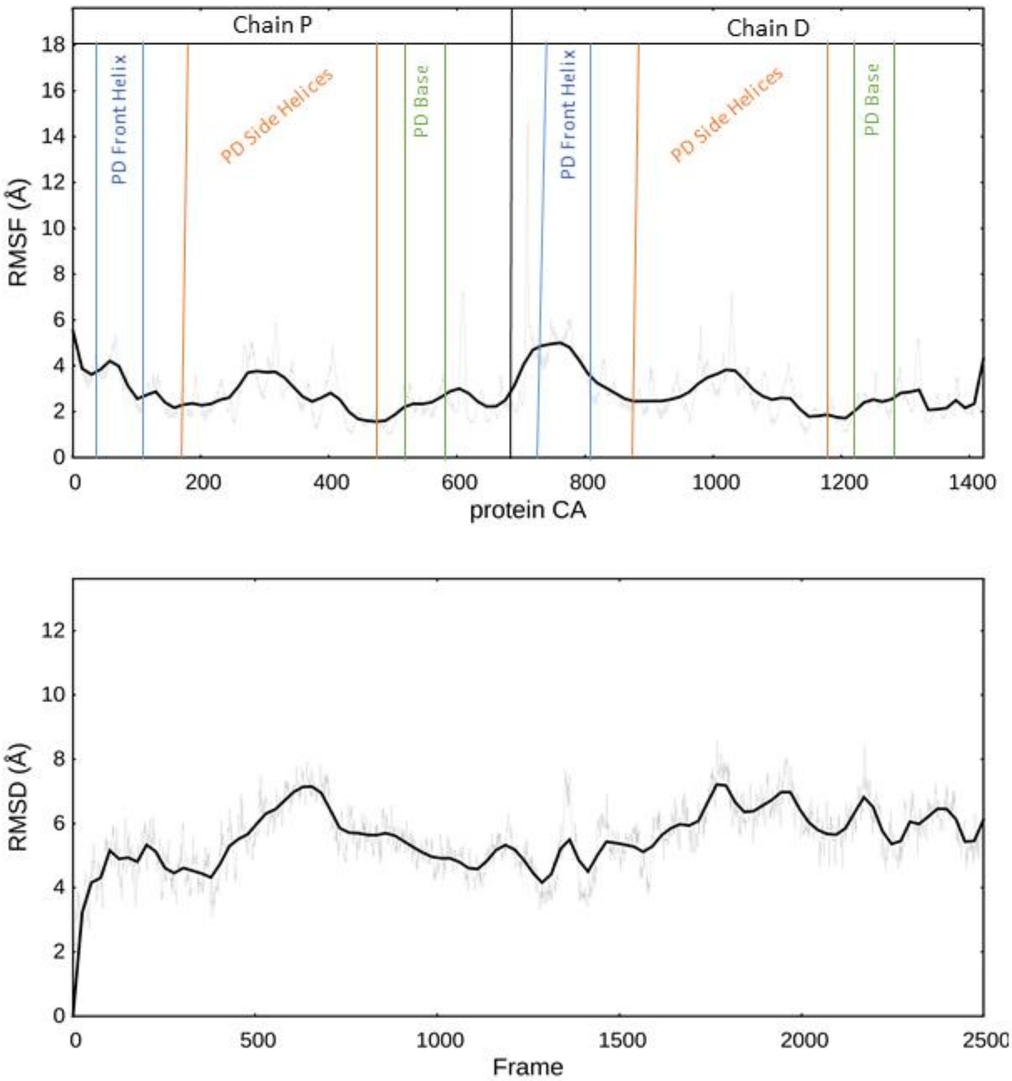


**Figure S5. ACE2 RMSF and RMSD during MixMD simulations in the presence of dHP. Results from MIXMD simulation.** ACE2 simulation indicated increased mobility in the extracellular domain, especially on the front helix on residues I21-H34, K74, on the external sides of the PD on residues N250, Y237, N338, F327 and the surrounding residues, indicating a moderate overall flexibility of the PD. The homodimer exhibited a symmetric behavior with the two “pincers” of the receptor showing the highest overall flexibility while the collar in proximity with the membrane remained more stable throughout the simulation. It is important to notice that the membrane was not included in the simulation and that the unity of the structure was kept by the strong intra-chain interactions alone.

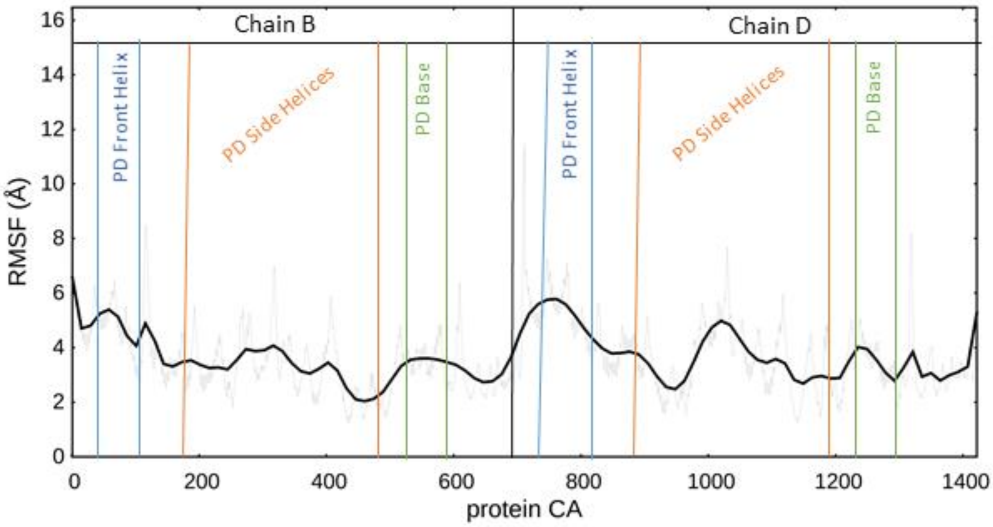


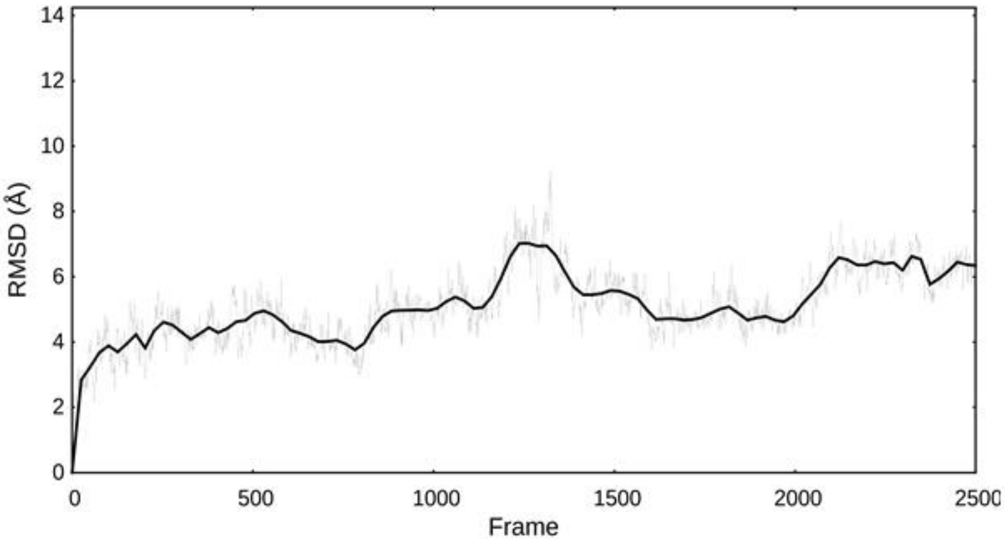


**Figure S6. ACE2 RMSF and RMSD during MixMD simulations in the presence of dHS. Results from MixMD simulation.**

In a similar scenario the overall flexibility of ACE2 PD was more accentuated on the external walls of the pinchers around residues E198, K596- S602, and on the helices on the ends of the pinchers on residues Q305-W328, R306-K313. The overall displacement of the structure fluctuated in presence of HS in respect to HP simulation, possibly due to the occurring of more frequent and durable contacts along the whole structure. HS clustered interactions at the base of ACE2 around residues W635, N636, R678, R705 is responsible for the increased mobility of the surrounding area which, consequentially, increased the movement of the whole structure, in absence of a stabilizing effect of the membrane. The more frequent contacts on the structure with HS, in respect of HP, allowed for a clearer definition of the density maps of the ligand along the structure.

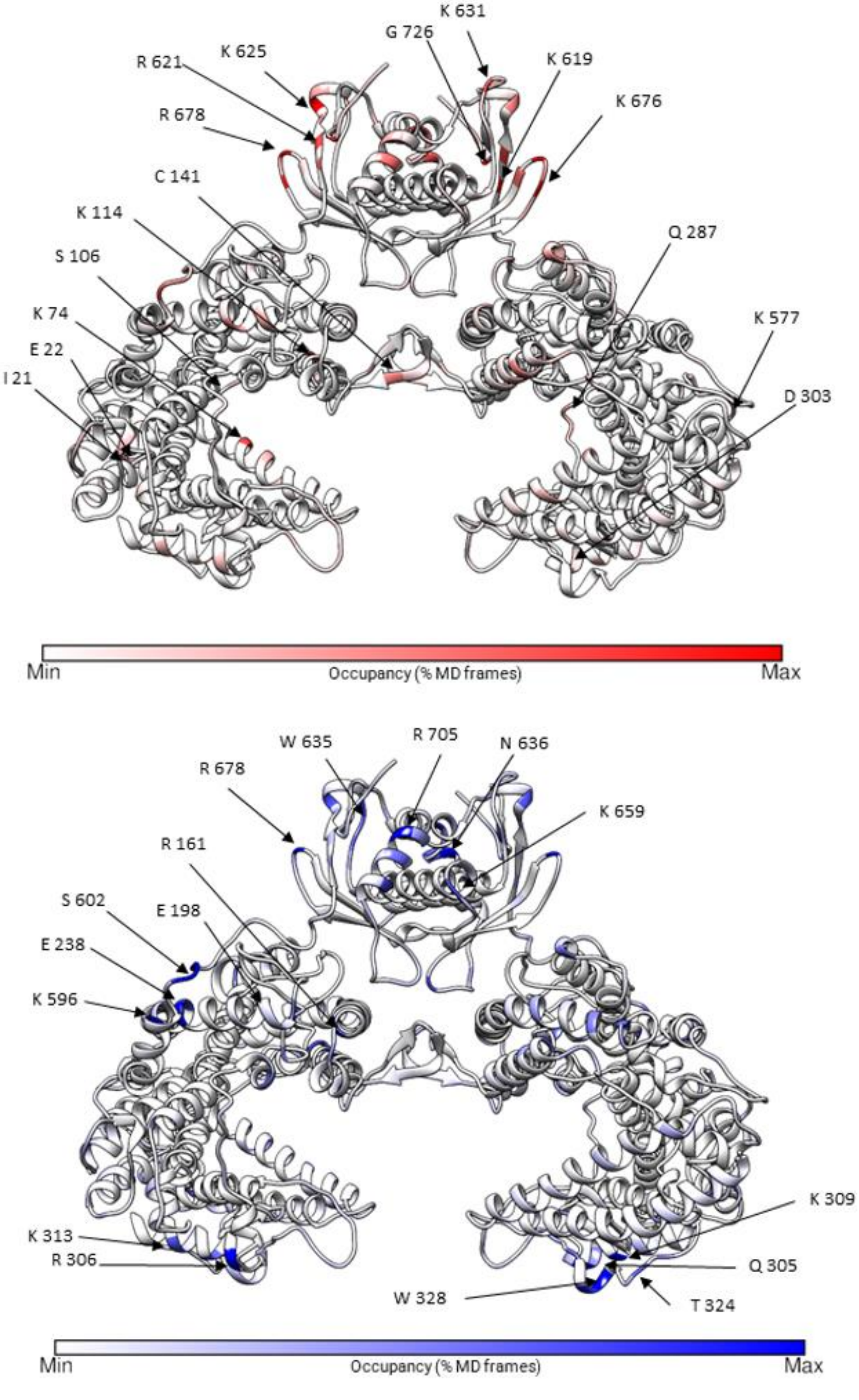


**Figure S7. S Contacts between ACE2 and dHP (red) and dHS (blue).**

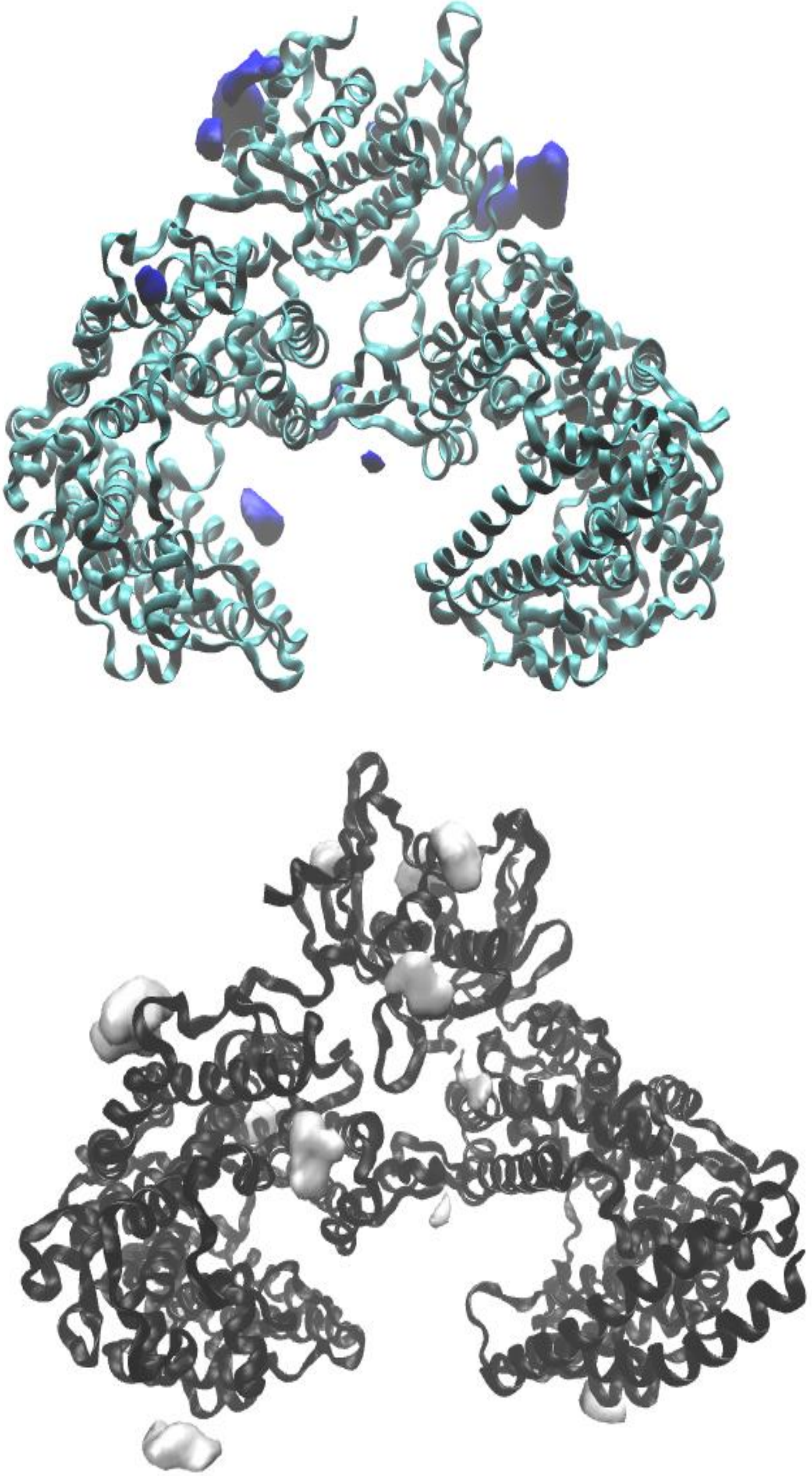

**Figure S8. Volumetric maps (ISOVALUE = 20%) of dHP (top) and dHS (bottom)**
